# NuclearIDTracker resolves intestinal cell identity and lineage dynamics through nuclear phenotypic signatures

**DOI:** 10.64898/2026.08.26.747251

**Authors:** Nguyen T.B. Nguyen, Rutger N.U. Kok, Sira Gevers, Xuan Zheng, Max A. Betjes, Lukas Ritter, Danny Feijtel, Matthew B. Smith, Sam F.B. van Beuningen, Jeroen S. van Zon, Sander J. Tans, María J. Rodríguez Colman

**Affiliations:** Molecular Cancer Research, Center for Molecular Medicine, University Medical Center Utrecht, Heidelberglaan 100, 3584 CG Utrecht, the Netherlands; Oncode Institute, The Netherlands; Hubrecht Institute, Royal Netherlands Academy of Arts and Sciences, University Medical Center Utrecht, the Netherlands; Autonomous Matter Department, AMOLF institute, Amsterdam, the Netherlands; Medicine, University of Basel; Lab of Cellular Disease Models, Department of Paediatrics, Regenerative Medicine Centre Utrecht, UMC Utrecht, Uppsalalaan 8, 3584 CT Utrecht, The Netherlands; Institute of Biology Leiden, Leiden University, Leiden, The Netherlands; Bionanoscience Department and Kavli Institute of Nanoscience Delft, Delft University of Technology, Delft, Netherlands; Cell and Chemical Biology, Leiden University Medical Center

## Abstract

Organoid models have transformed our understanding of intestinal renewal. Fluorescent imaging has been extensively used to identify key cell types and their differentiation pathways, but immunofluorescence provides only static readouts, whereas live imaging requires fluorescent-reporter engineering and is constrained by limited multiplexing and spectral overlap. Here, we introduce NuclearIDTracker, an explainable machine-learning framework that infers cell identity directly from 3D nuclear segmentations. Using a single nuclear marker, NuclearIDTracker accurately classifies intestinal cell types and integrates with single-cell tracking to resolve lineages and reconstruct dynamic state transitions during organoid development. We show that TA-like cells, rather than stem cells, drive early crypt formation and generate enterocyte and Paneth lineages, as well as the stem-cell population, which emerges only later and subsequently replenishes the TA-like compartment. Following stem-cell ablation, crypt regeneration was not driven by a single discrete cell type. Instead, multiple epithelial populations converged on a proliferative regenerative state with a nuclear phenotypic signature that resembled, but remained distinct from, that of homeostatic TA-like cells, and a YAP/TAZ-associated fetal-like transcriptional signature. Thus, nuclear phenotypic signatures resolve cell identity and reveal coordinated epithelial plasticity during crypt regeneration. NuclearIDTracker establishes a non-perturbative tool to quantify cell identity and state dynamics at single-cell resolution, revealing previously inaccessible biological dynamics and expanding the toolkit for studying epithelial homeostasis, regeneration, and disease.

## Introduction

The mammalian small intestine is a classical model of a rapidly self-renewing tissue, and intestinal organoids provide a powerful *in vitro* system that captures the major epithelial cell types and faithfully recapitulates key dynamics of epithelial homeostasis and regeneration^1,2^. Recently developed single-cell tracking approaches of time-lapse live imaging of organoids enable the study of cellular dynamics, including cell movement, division, death, and lineage reconstruction^3,4^. We recently extended this methodology by incorporating signal quantification from fluorescent reporters to infer cell identity and metabolic state during live imaging of tumor organoid development^5^. However, simultaneously mapping all cell types and their transitions during development remains challenging. Although several approaches can establish cell identity, including immunofluorescence^6^, smFISH^7^, and single-cell sequencing^8^, these methods provide static observations and do not resolve temporal dynamics^6^. Alternatively, genetically encoded fluorescent proteins controlled by cell-type-specific promoters can label distinct cell types and are compatible with live imaging, but their use is constrained by their availability, spectral overlap during multiplexing, and phototoxicity^9,10^. We therefore sought a minimally invasive and labor-efficient strategy to infer cell identity directly from existing imaging data.

Historically, cell identity has often been inferred from morphological features, and cell types were named after their microscopic appearance (e.g. rod, cone, or goblet cells)^11^. Recent advances in digital image processing have enabled automated classification of cellular phenotypes^12,13^, and inference of cellular trajectories and metastable states^14–17^. These studies suggest that cellular morphology can provide a non-invasive proxy for cellular identity, and state. However, most approaches have been applied to 2D models or to 3D models at discrete time points, and typically rely on whole-cell segmentation, which is challenging and error-prone in dense 3D epithelial models. Hence, methods for marker-free dynamic type identification in organoid models are currently lacking.

Cell identity is largely governed by gene expression programs, which are regulated not only by transcription factor activity but also by nuclear architecture and chromatin accessibility^18^, suggesting that nuclear morphology may reflect underlying cellular identity. Indeed, static 2D image analyses have shown that nuclear shape can predict cell type in human kidney and rat brain^19,20^. Advances in convolutional neural networks have considerably improved both nucleus and whole-cell segmentation^21–23^, with higher accuracy achieved for nuclear segmentation^24,25^. Nuclear morphometrics may therefore provide a robust and scalable strategy to infer cell identity in live 3D organoid systems. However, whether this approach can resolve intestinal epithelial cell types and lineage dynamics remains unknown.

Here, we present NuclearIDTracker, an explainable framework that infers cell identity and reconstructs identity transitions over time from nuclear morphometrics extracted from 3D live imaging. Applied to intestinal organoids, NuclearIDTracker classifies epithelial cell types and resolves dynamic state changes during de novo crypt formation and regeneration following stem-cell depletion. By decoding identity directly from nuclear phenotypic features, this approach reveals coordinated epithelial plasticity and establishes the nuclear phenotype as an intrinsic, non-perturbative readout of cell state in developing and regenerating epithelia.

## Results

### Nuclear phenotypic features identify intestinal cell types

To investigate whether nuclear features can distinguish cell types, we used mouse intestinal organoids genetically engineered to express an H2B-fused fluorescent protein to label nuclei. We performed live imaging of organoids followed by fixation and immunostaining to define specific cell types^6^ (**Figure 1A**). Immunostaining for olfactomedin 4 (OLFM4), lysozyme (LYZ), wheat germ agglutinin (WGA), keratin 20 (KRT20), and aldolase B (ALDOB) identified stem cells, Paneth cells, goblet cells, and enterocyte populations, respectively (**Figure 1C**). Goblet cells were rarely detected in control cultures; therefore, IL-4 and IL-13 were added to the culture medium to promote goblet cell differentiation (**Figure 1B**).

**Figure 1.**
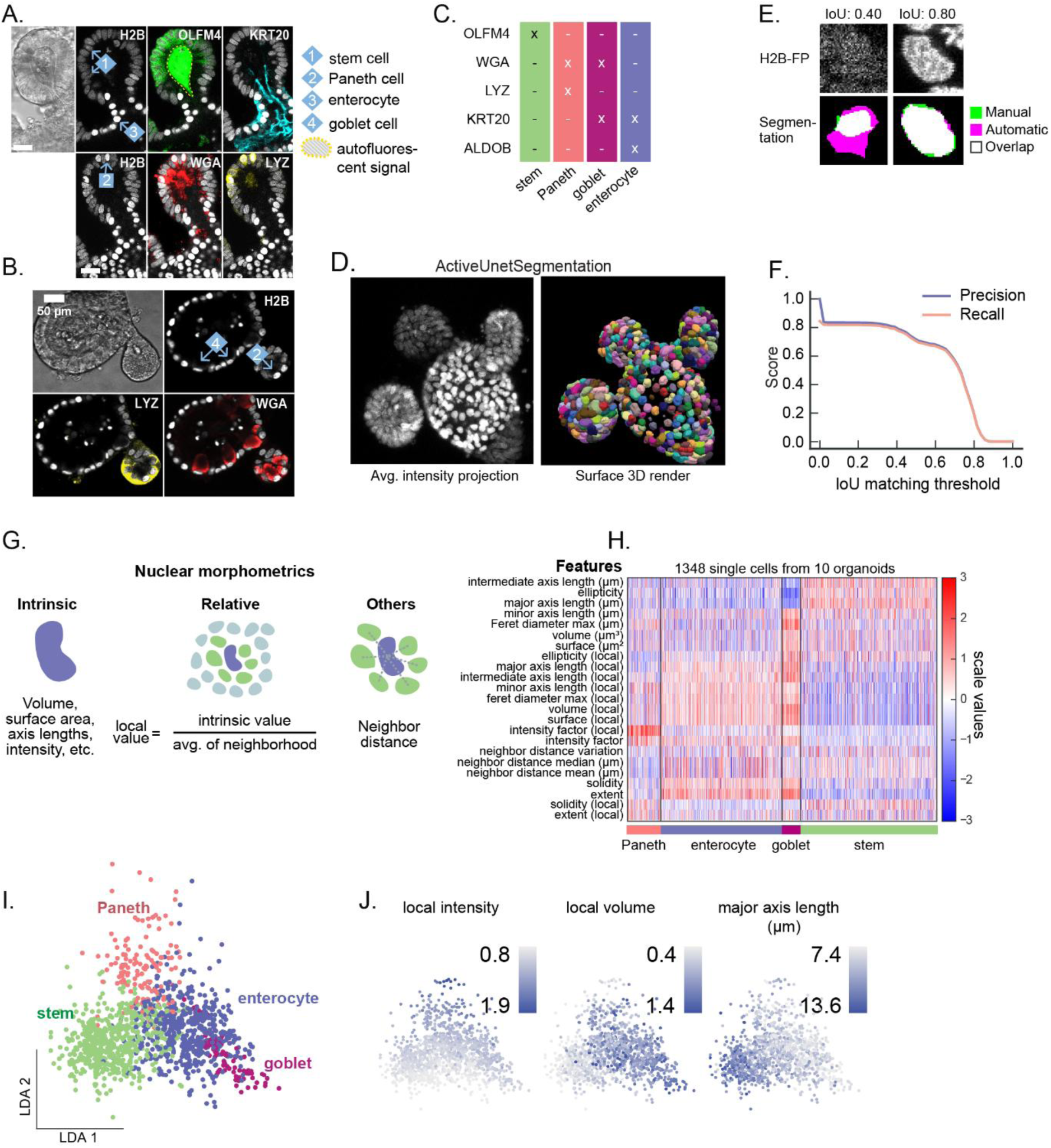
3D nuclear features separate intestinal cell types. **(A**) Examples of immunostaining on an intestinal crypt (scale bar = 35 µm). **(B)** Examples of immunostaining on a goblet cell-enriched (IL-4/IL-13-treated) organoid (scale bar = 50 µm). **(C)** Mapping of (immuno)staining to cell type. **(D)** Example of nucleus segmentation using ActiveUnetSegmentation. **(E)** Example of a bad and good segmentation with Intersection-over-Union (IoU) = 0.40 and 0.80, respectively. **(F)** Segmentation precision and recall compared to manual segmentation as a function of IoU matching thresholds. **(G)** Illustration showing the three different kinds of nuclear morphometric features: intrinsic, relative and others. **(H)** Heatmap of all features of all cells in the training set. Features were log-transformed, then scaled to a mean of 0 and standard deviation of 1. P-values were computed across organoids by using the average feature value of all cells within each organoid. **(I)** Linear Discriminant Analysis (LDA) plot of cell features, separated by cell types as determined by immunostaining. **(J)** The LDA-plot of panel I, colored by selected features.

For each stained organoid, we analyzed the final 15 time points prior to fixation. Nuclei were segmented in 3D using the H2B fluorescent signal and ActiveUnetSegmentation^25^, which employs a 3D U-Net neural network architecture to generate volumetric masks for individual nuclei (**Figure 1D**). Segmentation accuracy was assessed using the Intersection over Union (IoU) metric, which quantifies the overlap between manually annotated ground-truth masks and predicted segmentations. Example images of poor (IoU < 0.5) and good (IoU > 0.7) segmentations are shown in **Figure 1E**. The recall (fraction of true nuclei segmented correctly) and precision (fraction of predicted segmentations that are correct) curves indicated robust segmentation performance, with most segmented nuclei achieving an IoU > 0.7 (**Figure 1F**).

From the segmentation masks, we extracted 23 features, including nuclear shape descriptors (e.g., volume, surface area, and axis lengths), fluorescence intensity, local features capturing deviations from the surrounding microenvironment, and spatial-context metrics such as neighbor center-to-center distance (**Figure 1G**). Hereafter, we refer to this combined set as nuclear phenotypic features.

Next, we coupled nuclear phenotypic features to immunostaining-derived cell identities. Features from approximately 1300 nuclei were subjected to supervised dimensionality reduction using linear discriminant analysis (LDA), which revealed that these features separate distinct cell-type populations (**Figures 1H, I** and **Table S1**). Paneth cells were characterized by high local intensity; enterocytes and goblet cells by increased local volume; and stem cells by greater major-axis length (**Figure 1J**). These findings demonstrate that nuclear features generate distinct signatures associated with cell identity, supporting their use for predictive modeling.

### A logistic regression classifier infers cell identity

To predict cell types based on nuclear morphometrics, we developed a multinomial logistic regression model^26^. Unlike deep-learning approaches, this machine learning model is fully interpretable, allowing direct examination of which individual features contribute to cell-type prediction and how strongly they contribute. The model takes the phenotypic features of each nucleus as input and outputs a probability for each cell type (**Figure 2A**). Before being entered into the model, feature values were transformed to ensure comparability across parameters: ratio-based features were exponentially transformed to approximate a Gaussian distribution, and all variables were standardized to Z-scores (mean = 0, standard deviation = 1). The model was then trained using our dataset of nuclear features coupled to immunostaining-derived cell-type annotations. We hereafter refer to this framework as NuclearIDTracker.

**Figure 2.**
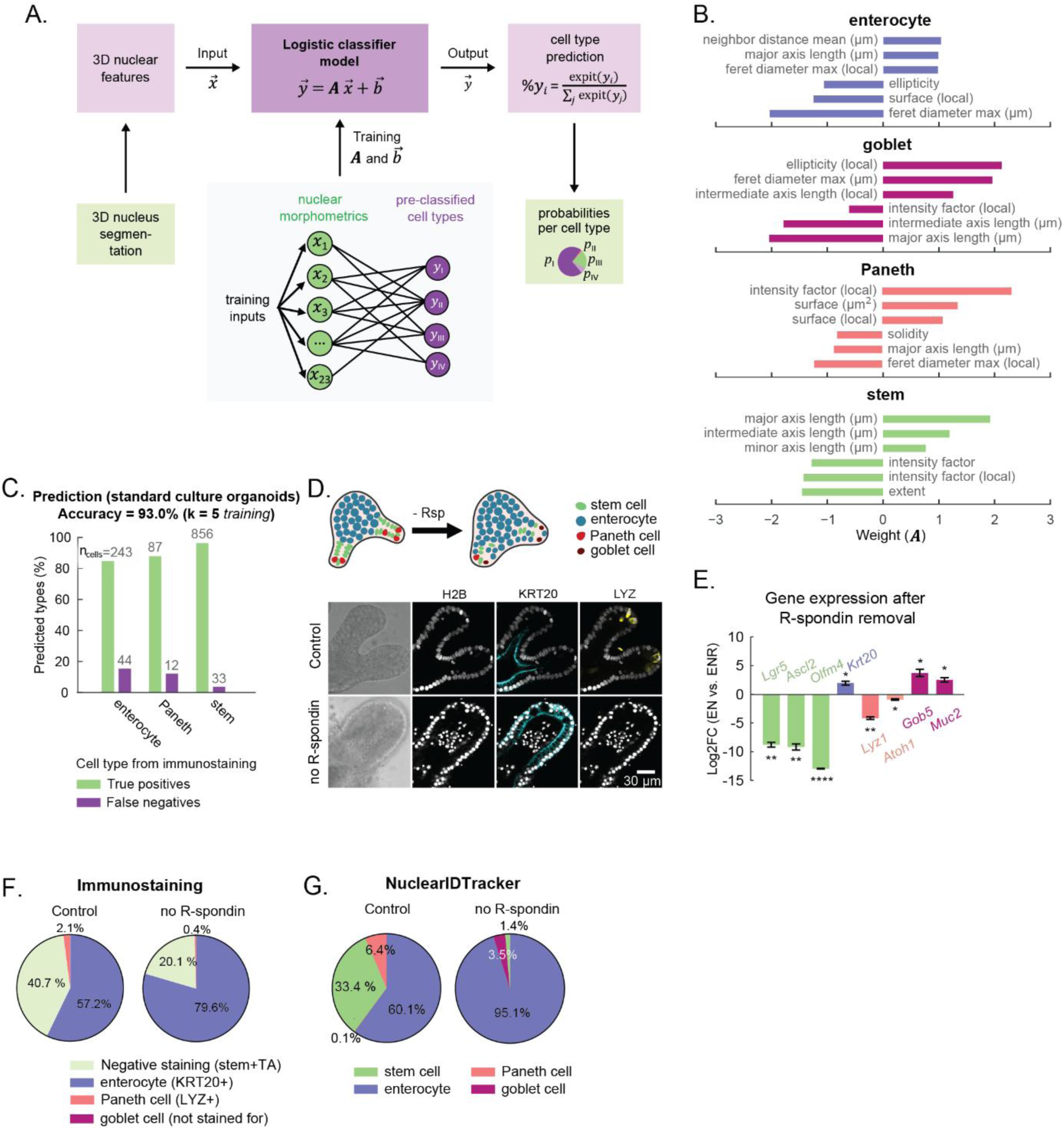
Cell type prediction using logistic regression classifier. **(A)** Illustration of the model. ***A*** is the matrix of the weights of all morphometric features for all cell types and *b⃗* a vector with the offset for each cell type. During training (fitting) of the model, ***A*** and *b⃗* are iteratively optimized to achieve maximum cell type prediction accuracy. *x⃗* are the measured features, and *y⃗* the resulting logit-likelihoods for each cell type. These are then normalized to obtain the likelihood for each cell type. **(B)** Top three most positive and most negative weights for each cell type. These features therefore contribute the most to the cell type classifcation. **(C)** True positives and false negatives for each cell type, measured for three control organoids. **(D)** Organoids grown without R-spondin for 72 hours. Immunostaining for KRT20 and lysozyme (LYZ) is shown for R-spondin-deprived and control organoids. Scale bar is 30 μm. **(E)** Differential expression of cell-type markers between control and R-spondin-deprived organoids (n = 3, log2FC, One sample t test, mean±SEM). **(F)** Immunofluorescence (cell percentage) for KRT20 or lysozyme (LYZ)-positive cells. Only successfully segmented nuclei were included (control: 3587 cells; R-spondin-deprived : 1071 cells). **(G)** Predicted cell-type fractions with and without R-spondin.

The relative contribution of each nuclear feature to cell-type classification was assessed by examining the trained model weights (coefficients). For each cell type, the six features with the largest positive and negative coefficients are shown in **Figure 2B**. For enterocytes, for instance, we found large positive weights for increased neighbor distances and major-axis lengths, suggesting that lower cell density and larger nuclei are predictive of this lineage. For Paneth cells, high nuclear intensity relative to neighboring nuclei emerged as the strongest positive predictor. In contrast, stem cells were characterized by low absolute and neighbor-relative nuclear intensity, together with large positive weights for major- and intermediate-axis lengths (**Figure 2B**).

We evaluated the performance of the NuclearIDTracker classifier using 5-fold cross-validation across a dataset comprising both standard-culture and cytokine-treated (IL-4/IL-13) organoids. The overall classification accuracy across all conditions was 85.4% (**Figure S2A**). Notably, performance was higher in the standard-culture organoid subset, reaching an overall accuracy of 93%, with OLFM4⁺ stem cells showing the highest accuracy (96.2%), followed by Paneth cells (87.9%) and enterocytes (84.7%) (**Figure 2C**).

Next, we tested our model under non-standard culture conditions by inducing enterocyte differentiation through R-spondin removal, thereby reducing Wnt signaling^27,28^. Gene expression analysis confirmed upregulation of the enterocyte marker *Krt20* and the downregulation of stem-cell markers (*Lgr5, Ascl2, Olfm4*) and Paneth-cell markers (*Lyz, Atoh1*) (**Figures 2D** and **2E**). Consistent with these transcriptional changes, immunostaining revealed an increase in KRT20⁺ enterocytes and a reduction in LYZ⁺ Paneth cells (**Figures 2D** and **2F**). NuclearIDTracker predictions indicated an increase in enterocytes and a marked decrease in stem and Paneth cells, mirroring the gene-expression and immunostaining results (**Figure 2G**).

Interestingly, NuclearIDTracker predicted a higher percentage of enterocytes than detected by KRT20 staining, with an approximately 15% discrepancy (**Figure 2G**). Further analysis showed that, in no-R-spondin organoids, predicted enterocytes displayed a distinct probability distribution, with fewer high-confidence predictions than under standard-culture conditions (**Figure S2H, S2I**). Thus, although enterocyte-like cells increased in abundance (**Figures 2F** and **2G**), not all may have reached a mature KRT20⁺ state. This suggests that NuclearIDTracker may identify early enterocyte-like states before the acquisition of detectable KRT20 expression. Consistent with this interpretation, KRT20-high cells showed higher enterocyte-probability scores than KRT20-low cells (**Figure S2I**). Furthermore, the NuclearIDTracker predicted a small population of goblet cells under this condition as well (**Figure 2G**). To validate this prediction, we assessed the expression of the goblet cell markers *Gob5* and *Muc2*, which supported an increase in goblet-cell differentiation following R-spondin removal (**Figure 2E**).

To assess how altered tissue architecture affects model performance, we treated organoid cultures with DAPT and CHIR99021, which inhibit Notch and activate Wnt signaling, respectively, inducing transient Paneth-cell enrichment at the expense of stem cells, followed by the loss of crypt integrity^28–30^. Although this treatment increased the abundance of LYZ⁺ Paneth cells, it also disrupted the normal interspersion of Paneth and stem cells (**Figures S2B** and **S2E**), thereby altering key local-context features used for Paneth-cell classification and causing NuclearIDTracker to underestimate Paneth-cell expansion (**Figures S2C–S2F**). Thus, Paneth-cell classification in strongly reorganized epithelia may require context-specific validation or retraining.

Together, these results establish NuclearIDTracker as a nuclear phenotypic framework for inferring intestinal cell identity and provide a foundation for evaluating its application to resolve dynamic cell-state transitions in time-lapse live-imaging datasets.

### NuclearIDTracker captures transitional cell states and lineage trajectories

NuclearIDTracker not only predicts the most likely cell type but also assigns a probability for every cell type. In the intestine, as stem cells migrate upward along the crypt-villus axis, they undergo progressive transitions through intermediate states before differentiating into mature enterocytes or secretory cell types, although these dynamics have not been directly visualized^6,27,32^. Thus, we asked whether NuclearIDTracker cell-type probabilities could reveal gradual cell-state transitions.

We first established a tri-color scale: green for stem cells, blue for enterocytes, and red for Paneth cells, with each color’s contribution proportional to predicted probability (**Figure 3A**). As goblet cells are rare under standard *in vitro* conditions, they were excluded from the scale. Mapping these probability-weighted colors onto cell positions along the crypt–lumen axis revealed the expected spatial organization^32^: predicted stem and Paneth cells localized to the crypt base, whereas predicted enterocytes were enriched in the villus-lumen region (**Figures 3B** and **3C**). Notably, cells located in the upper crypt (neck region) displayed a cyan coloration, indicating intermediate probabilities between stem and enterocyte identities (**Figure 3C**). This intermediate signature suggests that nuclear phenotypic features capture gradual cell state transitions during differentiation. Transit amplifying (TA) cells arise from Lgr5⁺ stem cells, retain high proliferative capacity, and serve as intermediate progenitors that generate enterocytes^32–34^. TA cells are typically positioned above the crypt base, along the stem-to-enterocyte trajectory near the crypt–villus interface. Interestingly, the spatial localization and intermediate probability profile of the cyan-marked cells therefore suggested that they may correspond to TA cells. We next asked whether NuclearIDTracker could identify TA cells directly from probabilistic outputs. To this end, we defined a stem-to-enterocyte (SE) axis, where a score of 1.0 represents 100% predicted stem identity and 0.0 represents 100% predicted enterocyte identity. Using a live Lgr5 reporter together with KI67 and KRT20 immunostaining, we mapped experimentally defined cell states onto this axis and observed a gradual shift in marker expression across SE scores from 0.55 to 0.30 (**Figure S3A**), consistent with progression from Lgr5⁺KI67⁺ stem cells to Lgr5^-^KI67⁺ cycling cells, toward KRT20⁺ enterocytes.

**Figure 3.**
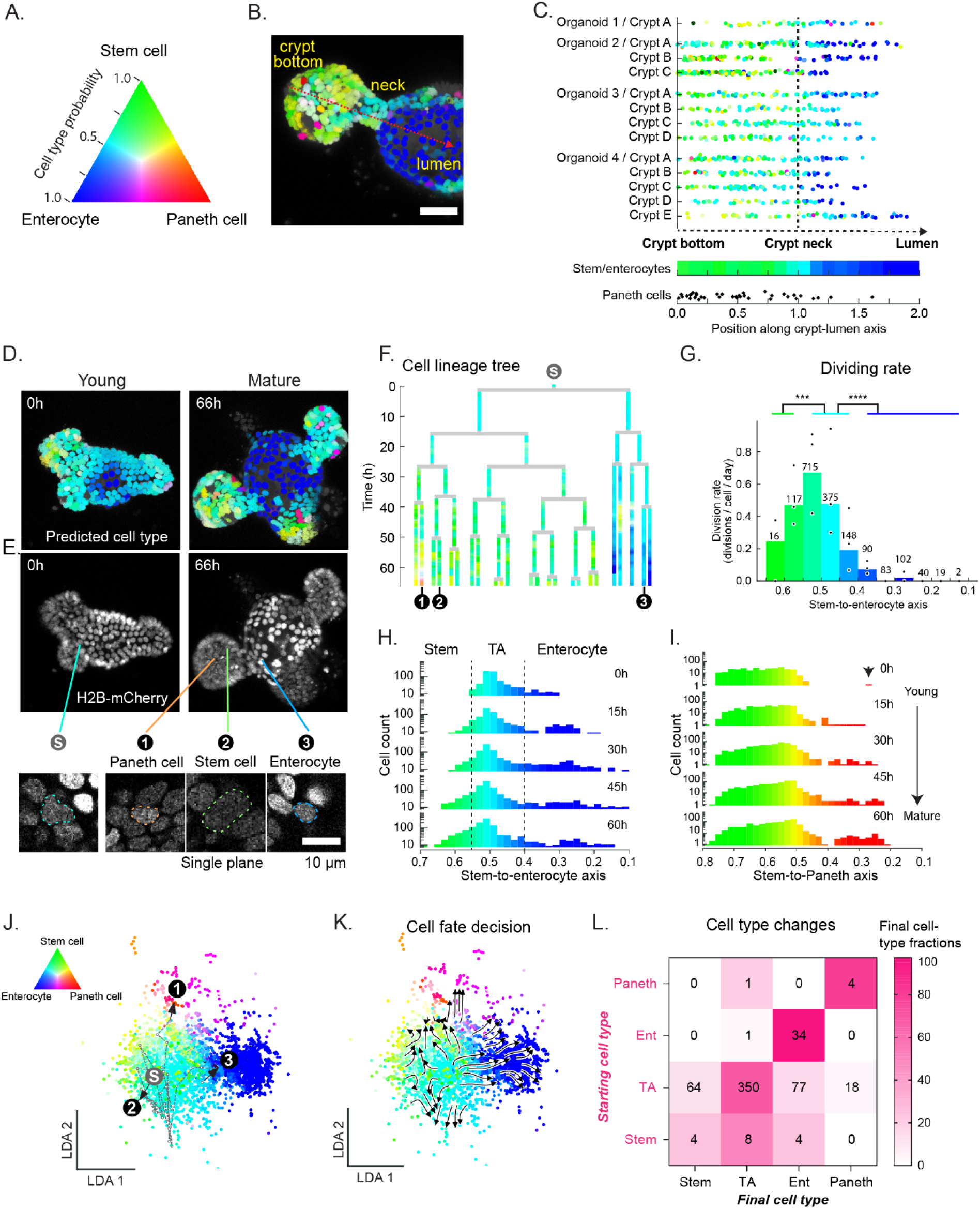
Predicting cell fate commitment using NuclearIDTracker. (**A**) Tri-color probability map of cell-type predictions computed by NuclearIDTracker. (**B**) A crypt from MSI organoids digitally colored by predicted cell type; the dashed arrow indicates the bottom crypt-neck-lumen axis. Scale bar 38.5 µm. (**C**) Distribution of cell-type probabilities (excluding Paneth cells) along the crypt-lumen axis across four organoids. (**D, E**) Time-lapse images of a developing organoid, digitally annotated by predicted cell type. Scale bar: 10 μm (**F**) Selected lineage tree of the organoid shown in panels (D,E). (**G**) Division rates of predicted cell types (excluding Paneth cells) plotted along the stem-to-enterocyte axis. Each point represents an individual organoid; bars indicate the average of three organoids. Number represent cell counts. Fisher’s exact test on pooled data of division counts and cell counts, ***: p<0.005, ****: p < 0.0001. (**H, I**) Cell counts over time along the stem-to-enterocyte and stem-to-Paneth axes for three organoids (logarithmic y-axis; see Methods section). (**J, K**) LDA plot of nuclear features from time-lapse data of all cells across three organoids, displaying both the overlaid example cell trajectories (see panel F) and the average trajectories of all cells at 5-hour intervals (see Methods). (**L**) Table showing counts of the predicted cell types at the start and end of 24-hour time-lapse videos from three organoids. Colors represent the fraction of each final cell type calculated per corresponding starting cell type. Only cells that were tracked for the entire duration were included. *Color intensity in (B, C, D, F-K) follows the scheme in (A)*.

Because a defining feature of TA cells is their proliferative capacity^33,34^, we next quantified division rates across cell states along the SE axis. To this end, we integrated NuclearIDTracker with OrganoidTracker^4,35^, enabling lineage reconstruction across successive divisions during organoid development (**Figures 3D-F**). This integration revealed gradual cell-state transitions in space and time, including stem-to-enterocyte progression within individual lineage trees, as illustrated by a representative lineage tracked throughout organoid development (**Figure 3F** and **Video S1**). <u>Not</u>ably, cells with intermediate SE scores showed the highest division rates, which declined significantly in cells with scores below 0.4 as they progressed toward terminal absorptive differentiation (**Figure 3G**). Importantly, this increased division rate was specific to intermediate SE scores along the stem-to-enterocyte trajectory and was absent along the stem-to-Paneth transition (**Figures 3G** and **S3C**). Together, these findings indicate that cells within the SE score range of 0.55–0.4, which are Lgr5^-^, correspond to TA-like cells (**Figure S3A**). In addition, no significant differences in cell cycle duration were observed among cells with SE scores between 0.4 and 0.6, with an average duration of 12 hours, consistent with previous reports for murine intestinal TA cells^33^ (**Figure S3B**). Complete cycles were rarely captured for high-probability stem cells (SE > 0.6), likely due to their lower frequency and suggesting longer cell-cycle durations relative to TA cells, as previously reported^33,36^. Together, these results define TA-like cells as a proliferative intermediate state along the stem-to-enterocyte trajectory, characterized by intermediate SE probabilities, high division rates, and progression toward absorptive differentiation.

### TA-like cells drive early organoid development and generate the stem-cell pool

Organoids derived from single cells initially grow as cystic structures, followed by the emergence of budding crypts^37^. To investigate cell-type composition and transitions during organoid development, we performed live imaging from early crypt formation through crypt maturation. NuclearIDTracker revealed progressive changes in cell-type composition over time (**Figures 3D**, **3H**, and **3I**). Mapping cells onto the SE axis showed that, at early stages, unexpectedly, no high-scoring stem cells were predicted, and most cells were TA-like, while a smaller fraction showed lower SE scores consistent with early enterocyte commitment (**Figure 3H**). Consistent with this observation, live Lgr5-GFP imaging combined with KI67 and KRT20 immunostaining demonstrated that early budding (young) organoids predominantly contained Lgr5⁻KI67⁺ TA cells and Lgr5⁻KRT20⁺ enterocytes (**Figures S3D** and **S3E**), confirming NuclearIDTracker predictions. High-scoring mature Paneth cells were predicted at early stages, (**Figure 3I**, 0h), and their abundance progressively increased over time. This was supported by staining of fixed organoids, which revealed an increase in the number of crypt-base WGA⁺ cells per organoid, from 12.33 ± 2.90 in young organoids to 22.00 ± 2.31 in mature organoids (**Figures 3I** and **S3E**). Notably, predicted stem cells with high scores emerged and expanded only later during crypt maturation, as confirmed by live imaging of Lgr5⁺ cells (**Figures 3H** and **S3D**).

To further investigate this finding, we analyzed cell-type annotations in published single-cell RNA-seq data from cystic and budding organoids following symmetry breaking^37^. This analysis confirmed that TA-like cells dominate early developmental stages, serving as a major proliferative reservoir during initial organoid expansion and subsequently giving rise to multiple lineages, including mature stem cells (**Figures S3F** and **S3G**).

To further explore the cell-state transitions, we tracked changes in predicted cell types over time. In **Figures 3D, E** and **F**, we highlighted a single starting cell (’**S**’) that divided into 24 progeny over 66 hours. Among these, three daughter cells (marked 1, 2, and 3) adopted distinct fates and occupied different spatial positions within the organoid (**Figure 3E**). The trajectory of nuclear feature changes during cell-type specification from the original ‘**S**’ cell to its stem-cell and differentiated progeny was visualized in a LDA landscape constructed from time-lapse data of three organoids (**Figure 3J**). Focusing on the initial 24 hours of growth, we computed average trajectories based on nuclear-feature changes at 5-hour intervals to generate a streamplot of all tracked cells (**Figure 3K**). Quantifying transitions based on initial and final predicted identities revealed that TA-like cells, which predominated at early stages, undergo self-renewal and generate stem cells, Paneth cells, and enterocytes (**Figures 3K** and **3L**). In contrast, mature enterocytes and Paneth cells largely retained their identities, with a low frequency of dedifferentiation events (**Figure 3L**). Although stem cells contributed to secretory and absorptive lineages, their their low abundance at early time points (**Figure 3L**). These findings show that TA-like cells drive early intestinal organoid development, acting as self-renewing progenitors that expand the stem-cell pool and generate secretory and absorptive lineages.

Together, these results show that NuclearIDTracker identifies mature and intermediate cell states in live organoids and, when integrated with single-cell tracking, reconstructs lineage trajectories revealing that TA-like cells proliferate, differentiate, and generate the later-emerging Lgr5⁺ stem-cell population during early organoid development.

### NuclearIDTracker reveals collective lineage convergence on a regenerative TA-like state

The intestinal epithelium displays remarkable self-renewal and regenerative capacity^38^. Recent studies have shown that, upon intestinal crypt damage, epithelial cells reprogram into a fetal-like state, driven by Wnt and YAP signaling, enabling proliferation-driven tissue repair^39,40^. Selective ablation of Lgr5^+^ stem cells has revealed that different Lgr5⁻ populations including Bmi1^+^ cells, Dll1^+^ secretory precursors and Alpi^+^ enterocyte precursors can replenish the stem cell pool^41–43^.

However, these studies largely rely on lineage tracing of one predefined cell population at a time, in which selected cell populations are genetically and fluorescently labeled to visualize their progeny at later time points. Although highly informative, this approach typically assesses the regenerative potential of individual populations rather than the collective and relative contributions of multiple epithelial cell states in real time. Thus, the collective behavior, relative contributions, and lineage preferences of different epithelial populations during crypt regeneration remain unresolved. Because NuclearIDTracker enables simultaneous cell-type identification and real-time visualization of cell-state transitions, it provides an approach to resolve collective lineage reorganization during intestinal regeneration.

To model stem cell loss, we used Lgr5-DTR-EGFP organoids and selectively ablated Lgr5^+^ stem cells with diphtheria toxin (DT), followed by DT washout to allow crypt regeneration (**Figure 4A**). As expected, DT treatment reduced total cell numbers and specifically the fraction of Lgr5^+^ stem cells, as indicated by live imaging of Lgr5-EGFP organoids and the reduced expression of stem cell-associated genes (*Lgr5, Olfm4, Ascl2, Ki67*) (**Figures 4B** and **S4A-C**). NuclearIDTracker predictions recapitulated the selective loss of cells with high stem cell probability following DT treatment (**Figure 4C**). In contrast, TA-like cells and enterocytes were minimally affected (**Figure 4C**). This prediction was supported by the presence of KI67⁺Lgr5⁻ survivor cells after treatment (**Figures 4B, S4B,** and **S4D**) and by the unchanged number of KRT20⁺ cells before and after DT exposure (**Figure S4D**). Consequently, the relative proportion, but not the absolute number, of KRT20⁺ enterocytes increased within the organoids (**Figures 4B** and **S4C**). The predicted Paneth-cell population also declined following stem cell ablation, consistent with reduced *Lyz1* expression and fewer crypt-based WGA^+^ cells in DT-treated organoids (**Figures 4D, S4B-E**).

**Figure 4.**
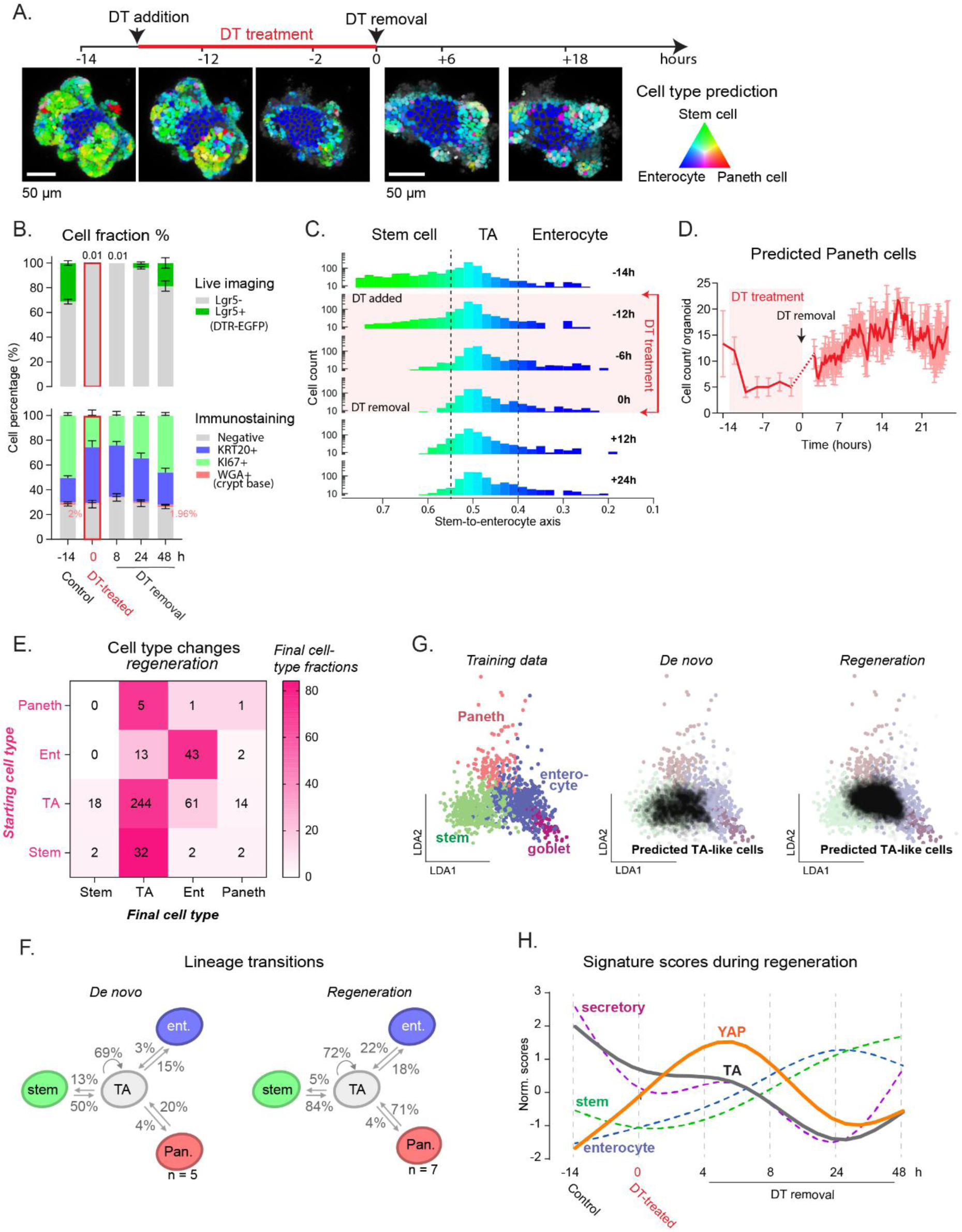
Cellular dynamics during crypt regeneration following stem cell ablation. **(A)** Maximum intensity projections of representative organoid, colored by predicted cell types, shown before, during, and after Lgr5⁺ stem cell ablation using diphtheria toxin (DT). **(B)** Quantification of Lgr5⁺ and Lgr5⁻ cell fractions per organoid (n = 3-4) using live imaging of Lgr5-DTR-EGFP organoids. Immunostaining of the same organoids was performed to assess KRT20⁺, KI67⁺, and WGA⁺ fractions before DT treatment, after 14 hours of DT exposure, and at 8, 24, and 48 hours post-DT washout. **(C)** Cell counts over time along the stem-to-enterocyte axis in three organoids (logarithmic y-axis; see Methods), captured before, during, and after DT-induced stem cell ablation. **(D)** Quantification of predicted Paneth cell numbers per organoid (mean±SEM, n = 3), before, during, and after DT treatment. **(E)** Table showing the counts of the predicted cell types at the start and end of 24-hour time-lapse videos of four regenerative organoids following DT washout. Only cells that were tracked for the entire duration were included. Dividing cells are represented twice (once per daughter). **(F)** Overview of lineage transitions between TA cells and other lineages (stem cells, enterocytes, Paneth cells) over the first 24 hours of development (from young budding organoids, n = 3) and regeneration (n = 4). Paneth cell counts are indicated. **(G)** Distribution of predicted TA-like cell positions on the LDA map (training data) during the final 8 hours of recorded videos of de novo (n = 3) and regenerative organoids (n = 4). **(H)** GSVA scores for signature gene sets (stem, TA, enterocyte, secretory and YAP; from Serra et al.^37^) calculated in mature organoids after 14 h of DT exposure and at 4, 8, 24 and 48 h following DT washout. Data were smoothed (spline fit, GraphPad Prism; 5 knots)) and z-scored per signature.

NuclearIDTracker revealed gradual recovery of stem cells during regeneration, which was supported by live-imaging analysis of Lgr5⁺-EGFP stem cells and transcriptional analysis of stem-cell markers (**Figures 4B**, **4C**, **S4B,** and **S4C**). Within the first 24 hours, the Lgr5⁺ cell fraction modestly increased from 0% to ∼4%, and became more prominent after 48 hours, reaching 19% (**Figure 4B**). In contrast, the Paneth cell population appeared to recover more rapidly, with proportions returning to pre-treatment levels after 48 hours (2.0% vs. 1.96%) (**Figures 4D** and **S4E**). This trend was also reflected in the Paneth-to-stem-cell ratio, which peaked 24 hours after DT removal and remained higher than pre-treatment levels at 48 hours (**Figures S4F** and **S4G**). Given the supportive role of Paneth cells in maintaining LGR5⁺ stem cells^31,44^, their earlier recovery may be essential for re-establishing the stem cell niche. The discrepancy between the predicted Paneth cells and the absolute number of WGA^+^ cells may reflect Paneth-cell precursors that are detected by the model before acquiring mature WGA expression (**Figures 4D** and **S4E**).

Time-lapse imaging of regenerating crypts during the first 24 hours revealed cell division rates comparable to those observed during de novo crypt development (**Figure S4H**). However, regeneration contained a higher fraction of cells with shorter cell cycles, potentially reflecting differences in cell-type composition and proliferative capacity **(Figure S4I**).

We next examined which cell types initiate and support crypt repair by tracking cell-state transitions during the first 24 hours of regeneration and comparing them with those observed during de novo development. A striking feature of regeneration was the pronounced expansion of the TA-like population (**Figures 4E** and **4F**). Specifically, enterocyte-to-TA transitions increased (from 3% to 22%) whereas Paneth-to-TA-like transitions increased from 20% to 71%, indicating that multiple differentiated populations converge on a TA-like state during early repair (**Figures 4E** and **4F**). TA-like cells largely retained their identity (∼70%) and showed similar rates of differentiation into enterocytes (15% vs. 18%) and Paneth cells (∼4%, **Figure 4F**). However, whereas TA-like cells contributed substantially to rebuilding the stem-cell pool during early development, their transition into stem-like cells was reduced during regeneration (13% vs. 5%), consistent with expansion of the TA-like compartment preceding stem-cell restoration (**Figure 4F**).

Interestingly, analysis of the LDA plots revealed that regenerative TA-like cells occupy a somewhat shifted position in nuclear phenotypic space compared with TA-like cells observed during de novo development (**Figure 4G**). Because regeneration has been associated with YAP-driven fetal-like reprogramming^39,40^, we next asked whether this regenerative TA-like state was transcriptionally aligned with fetal-like repair programs.

Transcriptomic analysis of regenerating organoids supported this interpretation. Following DT removal, YAP target genes were prominently upregulated, with distinct subsets peaking at different time points (**Figure S4J**). Gene set variation analysis (GSVA) revealed early and sustained enrichment of a YAP-activity signature during regeneration, whereas canonical stem-cell signatures decreased following DT treatment and were restored only at later stages (**Figures 4H** and **S4K**). Notably, the TA gene signature and TA marker genes were not induced during this early phase (**Figures 4H** and **S4K**), suggesting that the expanded TA-like population does not simply represent amplification of homeostatic TA cells. Rather, NuclearIDTracker identifies a proliferative intermediate state during regeneration that shares, but does not fully recapitulate, the nuclear phenotypic features of homeostatic TA cells and is transcriptionally aligned with fetal-like repair programs. Thus, regenerative TA-like cells are closest to homeostatic TA cells in nuclear phenotypic space, yet remain phenotypically and transcriptionally distinct.

Together, these data show that crypt regeneration after stem-cell ablation is characterized by the rapid expansion of a regenerative TA-like population with shorter cell cycles, the transient acquisition of YAP-associated fetal-like transcriptional features, and the delayed restoration of the Lgr5⁺ stem-cell compartment. NuclearIDTracker therefore reveals collective lineage reorganization during crypt regeneration, whereby multiple epithelial populations converge on a shared proliferative intermediate state during early repair.

## Discussion

In this study, we show that a single nuclear chromatin marker contains sufficient information to infer epithelial identity in live intestinal organoids. NuclearIDTracker turns the nuclear marker from a passive tracking label into an identity-resolving readout. This enables live reconstruction of epithelial plasticity and reveals that regeneration is not driven by a single reserve population, but by the convergence of multiple epithelial states onto a shared proliferative regenerative state.

A key strength of this approach is its probabilistic output, which enables the resolution of intermediate or transitional cellular states rather than enforcing discrete classifications. This is particularly relevant in dynamic epithelial systems, where lineage commitment is continuous and progressive rather than an on–off switch. Consistent with this, the model identifies intermediate TA cell states along the stem-to-enterocyte trajectory without pre-training on TA ground-truth labels. These findings indicate that nuclear phenotypic features capture information associated with lineage-specific transcriptional programs and may reflect chromatin reorganization during cell-fate transitions. This interpretation is supported by studies linking nuclear architecture and genome organization to cellular phenotype across diverse tissues^45–48^ and raises the possibility that nuclear phenotype is coupled to cell-fate regulation rather than representing a passive correlate. Determining how these nuclear changes relate to specific chromatin and transcriptional programs warrants further investigation.

The use of logistic regression provides an interpretable framework that identifies which nuclear features contribute to each cell-type prediction. This interpretability also helps explain changes in model performance under perturbation. For example, altered Paneth–stem cell organization following DAPT–CHIR treatment changed local-context features used for Paneth-cell classification. Thus, changes in model performance may reflect biologically relevant alterations in tissue organization and indicate when condition-specific validation or retraining is required.

Integrating NuclearIDTracker with OrganoidTracker^4,35^, enables reconstruction of cell-state trajectories across successive divisions. This provides access to division rates, cell-cycle duration, and lineage transitions. By relying on a single fluorescence channel, the method also limits phototoxicity and avoids the need for multiple cell-type-specific reporters, supporting long-term imaging in sensitive regenerative settings. Using the nuclear channel as an identity backbone further leaves additional channels available for functional reporters of signaling, metabolism, stress, or cell-cycle state.

During de novo organoid development, this approach revealed that TA-like cells dominate early crypt formation and generate enterocyte and Paneth lineages, as well as the later-emerging Lgr5⁺ stem-cell population. Once established, stem cells subsequently replenish the TA-like compartment. These findings indicate that early organoid development does not initially follow the conventional stem-to-progenitor hierarchy, but instead proceeds through a proliferative TA-like population that gives rise to the mature crypt lineages.

A central question in the field concerns the cellular origin of regeneration. Enterocyte and secretory progenitors, differentiated cells, surviving Lgr5⁺ stem cells, and Bmi1⁺ reserve populations have all been implicated^41,42,50,51^. Our high-spatiotemporal-resolution analysis suggests that early crypt regeneration proceeds through expansion of an Lgr5^low^ TA-like regenerative state rather than immediate restoration of the Lgr5⁺ stem-cell pool. This state arises from multiple epithelial sources, including surviving TA-like cells, enterocyte- and secretory-lineage cells, and remaining stem cells. These findings argue against a single fixed cellular origin and instead support a model in which several epithelial populations converge on a common proliferative state during early repair.

The regenerative TA-like population was closest to homeostatic TA cells in nuclear phenotypic space but remained phenotypically and transcriptionally distinct. During the first 24 hours following DT removal, these cells transiently acquired a YAP-associated fetal-like transcriptional program, consistent with previously described damage-induced regenerative states^52^. Similar Lgr5^low^ populations have been reported in independent studies^43,50,52^, although the extent to which these states overlap remains unresolved. Integrating NuclearIDTracker with signaling reporters could help define how such states emerge, interconvert, and return to homeostasis. Their cellular origins and dynamics are also likely to vary across injury models and stages of repair. Because our analysis focused on early regeneration, later transitions and the full return to homeostasis remain to be resolved.

Together, this work establishes nuclear phenotypic features as a live, non-perturbative readout of epithelial identity and provides a scalable strategy to reconstruct cell-state dynamics in organoids. Across both de novo development and regeneration, NuclearIDTracker reveals distinct uses of proliferative TA-like states: during development, they generate the later-emerging stem-cell population, whereas during regeneration, multiple epithelial populations converge on a YAP-associated regenerative state before restoration of the Lgr5⁺ compartment. NuclearIDTracker therefore provides a flexible platform for integrating long-term lineage tracking with live reporters of signaling, metabolism, and tissue repair.

### Limitations of the study

In this study, we prioritized the most common intestinal epithelial cell types, which are also the most abundant *in vivo* and in organoid systems. Therefore, model performance was not evaluated on rarer epithelial lineages, such as enteroendocrine, M, and tuft cells, which were absent from the training dataset. In addition, no specific marker for TA cells was included during training; TA states were therefore inferred indirectly as intermediate profiles between stem and enterocyte identities. The model employs standard logistic regression and therefore captures linear relationships between measured features and cell-type identity. While this reduces the risk of overfitting and improves interpretability, it may limit the ability to model nonlinear dependencies and feature interactions. Variation in classification accuracy across validated organoid sets indicates that epithelial integrity can influence nuclear morphology and, consequently, model performance (**Figure S2A**). We also assessed model performance on fixed organoids. Although accuracy decreased relative to live imaging, classification remained high (75.7%; **Figure S2G**). Retraining the model on fixed samples would likely further improve performance for endpoint applications. Finally, our approach depends on accurate nuclear segmentation and 3D digital reconstruction. Although reliance on nuclei rather than whole-cell boundaries facilitates implementation in dense 3D epithelial models, low segmentation accuracy may obscure morphometric differences between cell types and compromise classification fidelity. It should be noted, however, that rapid and continuous advances in 3D nuclear segmentation are likely to further improve robustness, accessibility, and transferability across imaging platforms.

## Data availability

Original code, algorithms, and computational models are available at: https://github.com/RodriguezColmanLab/NuclearIDTracker. RNA-sequencing data will be deposited in a public repository and made available upon publication.

## Acknowledgements

We would like to thank I. Verlaan (Snippert lab, UMC Utrecht) for preparing R-spondin- and Noggin-conditioned medium. We acknowledge the Utrecht Sequencing Facility (USEQ) for providing sequencing service and data. USEQ is subsidized by the University Medical Center Utrecht and The Netherlands X-omics Initiative (NWO project 184.034.019). This work was financially supported by VIDI VI.Vidi.203.008 financed by the Dutch Research Council (NWO), Stichting Proefdiervrij Nederland (https://proefdiervrij.nl/) and the Oncode Institute in the Netherlands. Work in the groups of S.J.T. and J.S.v.Z. are supported by the Netherlands Organization for Scientific Research (NWO). This work is supported by the project Organoids in Time with project no. 2019.085 of the research program NWO Investment Large financed by the Dutch Research Council (to J.S.v.Z. and S.J.T.)

## Contributions

Conceptualization, N.T.B.N., R.N.U.K., M.J.R.C.; methodology and investigation, R.N.U.K., N.T.B.N., S.G., L.R., M.A.B., J.S.v.Z, S.J.T. and M.J.R.C.; imaging acquisition, N.T.B.N, X.Z., M.A.B.; imaging analysis, R.N.U.K., N.T.B.N.; 3D segmentation strategy, M.B.S., S.F.B.B.; RNA sequencing, N.T.B.N., S.G.; manuscript writing and editing, N.T.B.N., R.N.U.K., S.J.T., M.J.R.C.; funding acquisition, D.F., J.S.v.Z., S.J.T, M.J.R.C.

---

**Figure S2.**
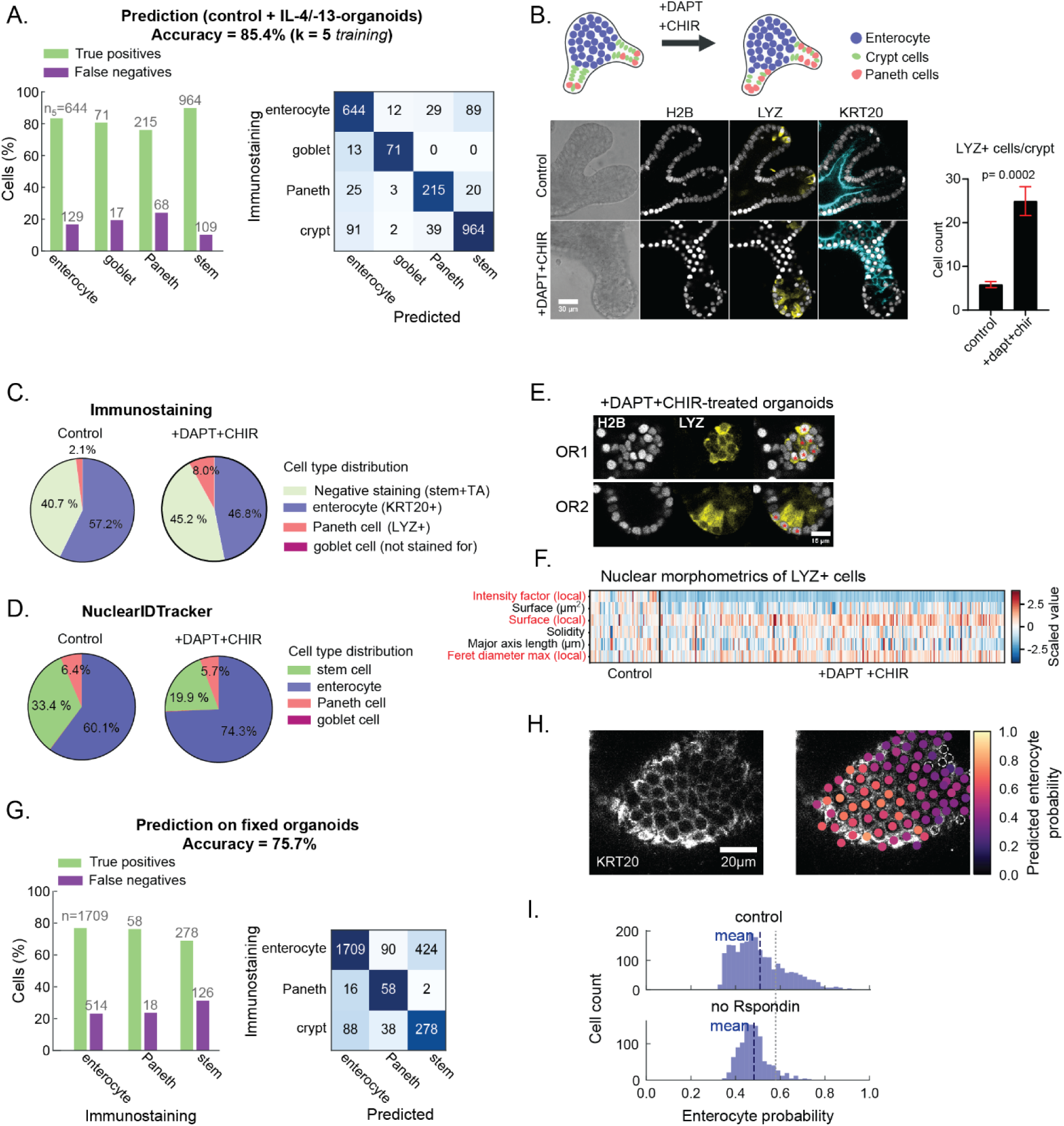
Performance of NuclearIDTracker across organoid conditions. **(A)** Comparison of predicted and immunostaining-derived cell types for both control and IL-4/IL-13-treated organoids from the training dataset (K=5-fold cross validation). **(B)** Immunostaining of DAPT + CHIR99021 (or CHIR)-treated organoids and controls with quantification of LYZ⁺ cells per crypt (n = 8 organoids per condition; mean±SEM; Mann-Whitney test). **(C)** Immunostaining for Lysozyme (LYZ) and KRT20 was quantified in control and DAPT+CHIR organoids and reported as the fraction of total nuclei (H2B-mScarlet) per condition (control: 3587 cells; DAPT+CHIR: 4397 cells). **(D)** Distribution of cell types, according to cell type prediction between control and DAPT+CHIR-treated organoids (n=8 organoids each). **(E)** Representative images of two crypts (OR1 and 2) showing LYZ⁺ cell localization at the crypt base in DAPT+CHIR-treated organoids (scale bar: 15 μm, asterisks indicate Paneth cells). **(F)** Heatmap illustrating changes in the top contributing features used to define Paneth cells (refer to Figure 2B) between control and DAPT+CHIR treatment. **(G)** Comparison of NuclearIDTracker-predicted cell types with immunostaining-based classifications (OLFM4⁺ stem cells, KRT20⁺ enterocytes, and LYZ⁺ Paneth cells) in fixed organoids. **(H)** Example of KRT20 staining along with predicted enterocyte probabilities. **(I)** Distribution of enterocyte probabilities in the control and R-spondin-deprived organoids.

**Figure S3.**
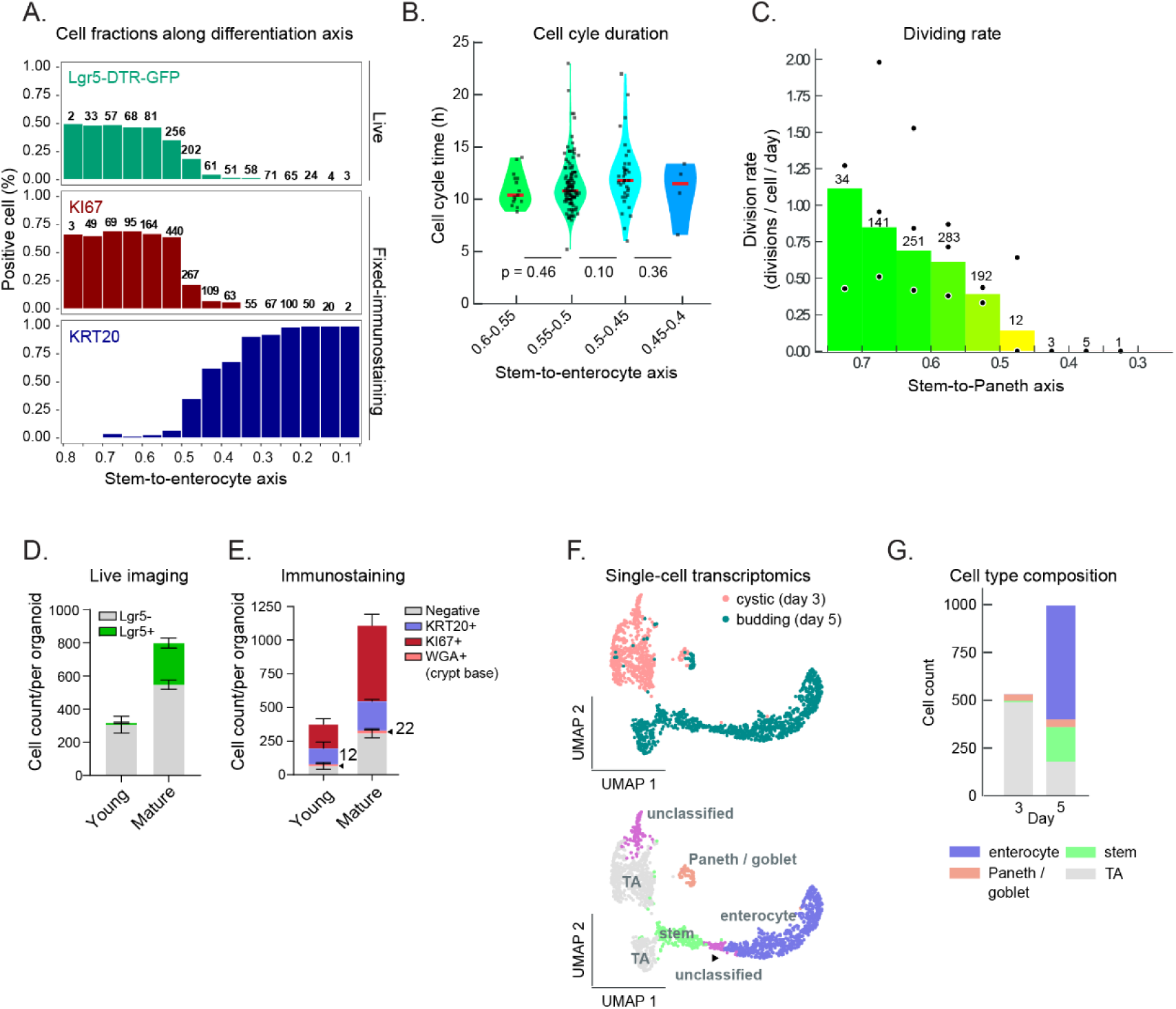
Predicting cell fate commitment using NuclearIDTracker. **(A)** Fraction of marker-positive cells; Lgr5 (via live imaging in MSI Lgr5-DTR-GFP organoids), KI67 and KRT20 (immunostained), plotted along the predicted stem-to-enterocyte axis. Both live imaging and immunostaining were performed on the same organoids. (B) Cell cycle length of all tracked cells from three organoids, displayed along the stem-to-enterocyte axis. Color intensity follows the scheme in Figure 3A. **(C)** Division rates of predicted cell types (excluding enterocytes) plotted along the stem-to-Paneth axis. Each point represents an individual organoid; bars indicate the average of three organoids. Numbers above the bars represent cell counts (Methods section). **(D,E)** Quantification of Lgr5⁺ and Lgr5⁻ cell numbers per organoid by live imaging of MSI Lgr5-DTR-GFP organoids. On the same organoids, immunostaining was performed to count cells positive for KRT20, KI67, and WGA, comparing young versus mature developmental stages. **(F)** UMAP plot of scRNAseq data from Serra et al.^37^, colored by the day after plating from single cell (top) and by annotated cell type (bottom). **(G)** Cell-type counts in the same scRNA-seq dataset as shown in (F, bottom panel). *Panels (A), (D-mature), and (E-mature) present the same dataset*.

**Figure S4.**
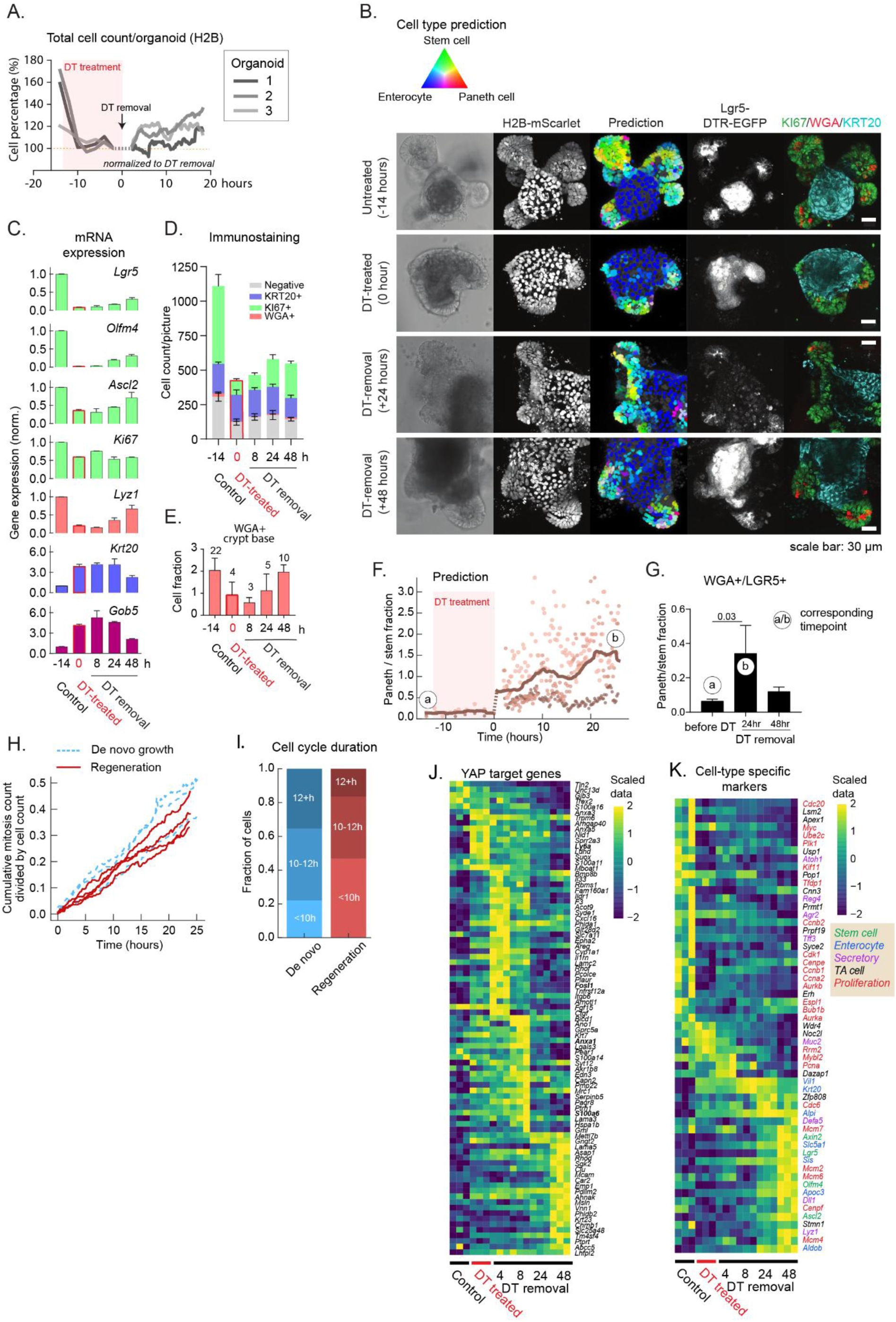
Cellular dynamics during crypt regeneration following stem cell ablation. **(A)** Quantification of total cell numbers (based on H2B-mScarlet) over time before, during, and after DT treatment. The imaging before and after DT removal was performed separately on the same organoids with different time resolutions (2 hours vs. 12 minutes) and z-stack coverage. Total cell counts were normalized to the timepoint closest to DT removal (0 hr). Data from three organoids are shown. **(B)** Representative images from live imaging (brightfield, H2B, Lgr5-DTR-EGFP) and corresponding immunostaining (KI67, WGA, KRT20) of untreated, DT-treated (14 hr), and regenerating organoids at 24 and 48 hours post-DT removal. Scale bar 30 µm. **(C)** Bulk mRNA expression of lineage markers in organoids before DT treatment, after 14 hours of DT, and at 8, 24, and 48 hours post-DT removal (n = 2 replicates, mean±SD). **(D,E)** Quantification of immunostained cells positive for KRT20, KI67, and crypt-based WGA in organoids before DT treatment, after 14 hours of DT, and at 8, 24, and 48 hours post-DT removal. WGA⁺ crypt-base fraction and total counts are displayed separately in (E) with an adjusted y-axis scale for visualization. Data represent n = 4 organoids (mean±SEM). **(F)** Ratio of predicted Paneth to stem cells over time, up to 24 hours post-DT removal. Data averaged from three organoids; timepoints [a] and [b] indicate pre-treatment and 24 hours post-washout, respectively. **(G)** WGA⁺ (crypt-based)/Lgr5⁺ (DTR-EGFP) cell fractions normalized to either fixed image or live-imaged total cells, before treatment ([a]), and at 24 ([b]) and 48 hours post DT-washout. **(H)** Cumulative number of mitotic events captured by live imaging over time during de novo growth or after DT washout (regeneration). **(I)** Fractions of cells with different cell-cycle durations, pooled from de novo (219 cells from 3 organoids) and regenerative (476 cells from 4 organoids) conditions. **(J,K)** Heat map of significantly expressed genes (FDR < 0.001), including YAP target genes and cell-type-specific genes, across different samples: control mature organoids, 14 h DT-treated organoids, and at 4, 8, 24 and 48 h following DT washout. Values were min-max normalized and rescaled to [-2,2] for visualization.

**Table S1.**
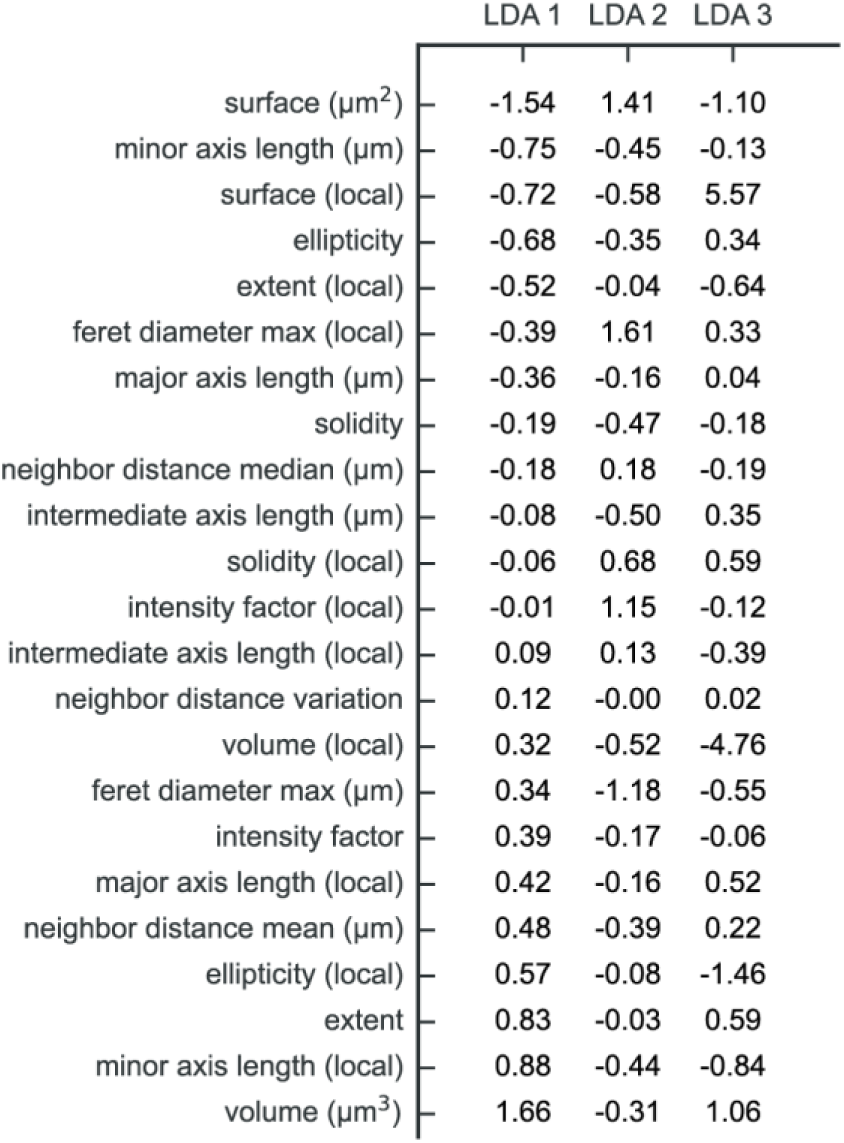
LDA scalings for each feature and LDA axis. Positive scalings indicate that the given feature increases along the given axis, and vice versa. Rows are sorted by the scalings for LDA 1.

**<u>Video S1</u>. NuclearIDTracker resolves cell identity and lineage transitions during intestinal organoid development.** Time-lapse imaging of an H2B-FP (Fluorescent protein: mCherry)-labeled intestinal organoid is shown together with NuclearIDTracker predictions, a highlighted example lineage, and the corresponding lineage reconstruction. Predicted cell identities are mapped onto individual nuclei over time, enabling simultaneous visualization of organoid growth, cell-state transitions, and lineage relationships.

## Methods and Materials

### Organoid lines

The mouse small intestinal (MSI) H2B-mCherry line was used in our previous publications^6,53^, and was originally a gift of Norman Sachs and Joep Beumer from the H. Clevers lab. For regeneration assays and immunostaining control validation, organoids derived from Lgr5^DTR-eGFP^ mice^43^ were transduced with a hEF1α-H2B-mScarlet-IRES-blasticidin cassette. This line was a gift from the labs of Gloerich (UMC Utrecht) and Sonnen (Hubrecht Institute, Utrecht).

For passaging, MSI H2B-mcherry and MSI Lgr5^DTR-eGFP^ H2B-mScarlet organoids were mechanically sheared into small clumps of cells and plated in Matrigel in based-ENR medium. The ENR medium contained DMEM F12 supplemented with 1% penicillin/streptomycin, 10mM HEPES, 2mM Glutamine, 1 × B27 (Life Technologies), 1.25 mM N-acetylcysteine (Sigma), murine recombinant epidermal growth factor (Peprotech), R-spondin1-CM (5% v/v, homemade) and noggin-CM (10% v/v, homemade).Organoids were kept at 37 °C and at 5% CO_2_.

To compare developmental stages, organoids were collected or fixed (PFA 4%, 15 minutes, room temperature) at two time points: onset of budding (young, 2-3 days post-shearing) and the mature crypt stage (5 days post-shearing).

### Training dataset construction

The training of our models used datasets reported in our prior study^6^, which provides detailed protocols for organoid culture, imaging, staining and cell tracking. Additional training data with goblet cell enrichment were generated using the same culture and preparation workflow^6^, with IL-4 and IL-13 added at imaging onset (10 ng/ml each, Peprotech). Organoids were imaged every 12 minutes for 61 hours using a Nikon A1R MP confocal microscope (40x oil-immersion objective, NA = 1.30, z-step = 2 μm) at 37°C/5% CO_2_. After imaging, organoids were immunostained for Lysozyme and WGA following the same protocol^6^. Cells were tracked automatically using OrganoidTracker^35^, and the trajectories of lysosome and WGA-positive cells were manually corrected. Only the final 3 hours of each time-lapse movie were included in the training dataset.

### Quantification of 3D nuclear feature parameters

Nuclei were segmented in each time point using ActiveUnetSegmentation^25^. Using skimage.measure.marching_cubes from scikit-image^54^ a mesh of each nucleus was reconstructed in micrometers coordinates. From this mesh, the following features were calculated: volume, solidity, surface, Feret diameter max, extent, minor axis length, intermediate axis length, major axis length and ellipticity. In addition, an ‘intensity factor’ was computed, defined as the mean intensity within each nuclear volume divided by the median intensity of all nuclei at the same time point. From each parameter, we calculated a local-normalized value by dividing the nucleus’s value by the mean of its six nearest neighbors. Finally, the mean, median and the median absolute deviation of the center-to-center distances to these six closest neighbors were recorded.

### Training and prediction procedure

For training, we used sklearn.linear_model.LogisticRegression from scikit-learn^55^, with per-nucleus feature vectors (all parameters described above) as inputs and four cell types defined by immunostaining as ground truth labels (**Figure 1C**). Local features were defined as the ratio between a nucleus’s feature value and the mean value of its six nearest neighbors. Ratio-type parameters, which are neighbor distance variation, solidity, ellipticity, the intensity factor, and all local metrics, were exponentiated. All parameters were then scaled linearly (z-scored; mean 0, SD 1) using sklearn.preprocessing.StandardScaler. The resulting scaling factors and offsets were kept. We performed 5-fold cross validation (*K = 5*), resulting in 5 models, of which one arbitrary model (the first one) was kept.

For cell-type prediction, features of the prediction data were standardized using the scaling factors determined from the training set, after which the trained model was applied to predict cell types.

To avoid training on truncated nuclei, we excluded any nucleus whose segmented mask touched the border of the 3D image volume. During cell type prediction, to avoid losing nuclei that slightly brush the border of the image, this step was omitted.

### Tracking cell-type trajectories over time

Single-cell linking from time-lapse data was obtained using OrganoidTracker^35^. OrganoidTracker-derived centroids were matched to segmentation masks to integrate tracking information with predicted cell types. A strict one-to-one correspondence between the two data were required: if either zero or more than one centroid fell within a nucleus mask, that mask was discarded for that frame. To reduce frame-to-frame noise, per-nucleus features were smoothed over time using a moving average window spanning ±1 hour.

### Linear Discriminant Analysis and stream plots

All nuclear features were scaled as described above. A Linear Discriminant Analysis (LDA) model was then fit into the training data using sklearn.discriminant_analysis.LinearDiscriminantAnalysis from scikit-learn^55^. **Figure 1I** shows the LDA projection of the training set only. In **Figures 3J**, **3K and 4G** that include cells not used for training, features were scaled using the mean and standard deviation *of the training data*, and the previously fitted LDA model was applied without re-fitting.

To construct the stream plot in **Figures 3J** and **3K**, LDA coordinates were calculated per cell and time point. The LDA coordinates were then downsampled by averaging within 5-hour bins (one mean per bin). For each bin, the starting LDA coordinates and changes (called deltas) in coordinates were recorded. The LDA plot was divided into a 25 x 25 square grid, ranging from the LDA coords (-5, 5) x (-5, 5). Every recorded delta was assigned to the grid cell based on its starting coordinates. For every grid cell, a stream value was defined: either the average of all deltas, or zero if less than two deltas were assigned to that cell. The resulting stream plot was plotted using Matplotlib’s^56^ Axes.streamplot function with a density parameter value of 0.9.

### Secretory/enterocyte lineage perturbation and enrichment

Following 3 days in ENR, or upon crypt formation, treatments were applied for 72 hours: CHIR99021 (3 µM) together with DAPT (10 µM), or EN medium (R-spondin1 removed). Organoids were then fixed in 4% PFA for 15 min at room temperature (RT), washed in PBS, and stored in PBS until staining. Untreated and EN conditions were harvested in parallel for RNA extraction.

### Organoid regeneration after DT treatment

MSI LGR5^DTR-eGFP^ H2B-mScarlet organoids were grown in ENR to the mature crypt stage (5 days post-shearing). Diphtheria toxin (DT, 30 ng/ml) was applied for 14 hours. DT was then removed by four sequential medium washes with 15-min incubations between washes, after which ENR medium was readded.

For time-lapse microscopy, organoids were imaged immediately after DT addition at 2-h intervals, recording only the H2B-mScarlet channel (excitation 570 nm, emission 590–675 nm) on a Leica SP8 confocal microscope (40× objective, z-step 2 µm). After DT washout, organoids were returned to the microscope 2-3 hours later for the regeneration phase. Imaging of the same organoids continued for an additional 24 hours at 12-minute intervals.

For static imaging and fixation, live imaging of Lgr5^DTR-eGFP^ signal was performed at 5 time points: before DT, after 14-hour DT, and at 8, 24, and 48 hour post-DT removal. Following live imaging, organoids were fixed within 30 minutes in 4% PFA for 15 min at RT.

RNA samples were collected at the same time points as the live/static imaging series (pre-DT, 14 h DT, and 8/24/48 h post-washout).

### Immunofluorescent (IF) staining

Mechanically sheared organoids were plated in precooled WillCo dishes (HBST-3522). After treatment or live imaging, wells were washed with ice-cold PBS. Next, they were fixed by 4% PFA for 15 minutes at RT and stored in PBS at 4 °C for up to 4 days. For staining, organoids were permeabilized/blocked with PBS buffer containing 10% DMSO, 2% Triton X-100, and 10 g/L BSA for 2 hours at 4 °C. Organoids were stained overnight with primary antibodies (KRT20 1:200, lysozyme 1:500, KI67 1:200) followed by Alexa fluorophore-conjugated secondary antibodies (+DAPI and +WGA 1:2000) for 4 hours at 4 °C. Imaging was performed using a SP8 confocal microscope (Leica).

### RNA extraction and real-time PCR

Organoid cultures subjected to different treatments were collected in ice-cold PBS. RNA purification was performed with the NucleoSpin RNA, Mini kit for RNA purification with DNase treatment (Macherey-nagel), following the manufacturer’s protocol. DNase treated RNA was used for cDNA synthesis using iScript cDNA synthesis kit (Biorad). Afterward, cDNA was subjected to qPCR using PowerTrack™ SYBR Green Master Mix for qPCR (Applied Biosystems™). Relative gene expression was quantified by the 2^−ΔΔCt^ method. Ct values were normalized to housekeeping genes (*Hnrnpa1, Gapdh, and β-Actin*). For ENR vs EN comparisons, only *Gapdh* and *β-Actin* were used as housekeeping genes. Reference conditions (untreated, ENR organoids, or young-stage organoids) were chosen as appropriate for each comparison. Primer sequences:

*qmHNRNPA1_fw_TGACAGCTATAACAACGGAG, qmHNRNPA1_rv_ AAAGTTTCCTCCCTTCATCG,*

*qmGapdh_fw_GAGAAACCTGCCAAGTATGA, qmGapdh_rv_CTCAGTGTAGCCCAAGATG,*

*qmbActin_fw_CTGTCGAGTCGCGTCCA, qmbActin_rv_TCATCCATGGCGAACTGGTG,*

*qmLyz1_fw_CGTTGTGAGTTGGCCAGAA, qmLyz1_rv_GCTAAACACACCCAGTCAGC,*

*qmGob5_fw_ACTAAGGTGGCCTACCTCCAA, qmGob5_rv_GGAGGTGACAGTCAAGGTGAGA,*

*qmMuc2_fw_ATGCCCACCTCCTCAAAGAC, qmMuc2_rv_GTAGTTTCCGTTGGAACAGTGAA,*

*qmAtoh1_fw_GCCTTGCCGGACTCGCTTCTC, qmAtoh1_rv_TCTGTGCCATCATCGCTGTTAGGG,*

*qmLgr5_fw_ GTTCAAGATGAGCGGGACCT, qmLgr5_rv_ ATAGGTGCTCACAGGGCTTG,*

*qmAscl2_fw_ CGTGAAGCTGGTGAACTTGG, qmAscl2_rv_ GGATGTACTCCACGGCTGAG,*

*qmOlfm4_fw_ TGAAGGAGATGCAAAAACTGG, qmOlfm4_rv_ CTCCAGCTTCTCTACCAAGAGG,*

*qmKrt20_fw_ TCGAGGTCCAAGTCACGGAG, qmKrt20_rv_ GCTCCAGAGACTCTTTCATGCT,*

*qmKi67_fw_ CCTTTGCTGTCCCCGAAGA, qmKi67_rv_ GGCTTCTCATCTGTTGCTTCCT*.

### Projection of single cells onto stem-to-enterocyte/Paneth axes

The stem-to-enterocyte (SE) axis location for each cell was computed as follows:

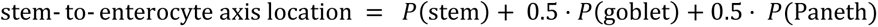

Here, *P*(*x*) is the predicted probability for cell type *x* for the cell of interest. Because *P*(stem) + *P*(goblet) + *P*(Paneth) + *P*(enterocyte) = 1, one can show that this formula is equal to 1 − (*P*(enterocyte) + 0.5 ⋅ *P*(goblet) + 0.5 ⋅ *P*(Paneth)). SE locations were calculated only for cells predicted as stem or enterocyte; cells predicted as Paneth or goblet were excluded.

For the stem-to-Paneth axis, the same procedure was followed: *P*(stem) + 0.5 ⋅ *P*(goblet) + 0.5 ⋅ *P*(enterocyte)

### Division rate measurement

We considered all cell tracks that are at least 2 hours long. A cell track is a trajectory of a single cell from its first detected frame (cell birth, start of time lapse, or cell entered imaged volume) to its last observable frame (mitosis, cell death, end of time lapse, or cell left imaged volume). For each track, a single cell type or location in the stem-to-enterocyte/Paneth axis was assigned. Since cell-type probabilities can change during the cell’s lifetime, we used the time-averaged per-frame cell-type probabilities across the entire track. The cell division rate is then calculated as the number of cell divisions, divided by the summed length of all tracks of the cell type.

Cell counts (above the bars in the bar plots) are the eventual cell counts after all divisions. For example, if a cell divides, and both daughter cells divide again, the cell count would be 4.

### scRNAseq analysis

scRNA-seq datasets from cystic and budding MSI organoids (C57BL/6 wild type; day 3 and day 5 post single-cell trypsinization) were obtained from GEO: GSE115956^40^. ENSEMBL gene IDs were converted to gene symbols using gprofiler2 (v0.2.3)^57^, after which duplicate genes were removed by retaining the entry with the higher count. Raw UMI matrices were loaded into a Seurat object (v4.3.1), and cells were filtered to retain those with 200–6,000 detected features and <15% mitochondrial transcripts. Data were then normalized and scaled using Seurat’s standard workflow^58^. Cell types were assigned with scType^59^ using marker sets adapted from Serra et al.^37^. Marker panels included: stem (*Lgr5, Olfm4, Ascl2, Axin2*), TA (20 TA markers from Serra et al.^37^), enterocyte (*Krt20, AlpI, Slc5A1, Vil1, Sis, Aldob, Apoc3*), and Paneth/Goblet (*Reg4, Atoh1, Dll1, Lyz1, Defa5, Muc2, Tff3, Gob5, Agr2*).

### RNAseq analysis

MSI Lgr5-DTR-EGFP H2B-mScarlet organoids were cultured in ENR medium and collected at two developmental stages: the onset of budding (young organoids, 2 days post-shearing) and the mature crypt stage (5 days post-shearing). To study transcriptional dynamics during regeneration, mature organoids (5 days post-shearing) were treated with DT (30 ng/mL) for 14 hours. DT was subsequently removed by four sequential medium washes, each followed by a 15-minute incubation, after which fresh ENR medium was re-added. Organoids were harvested at 0, 4, 8, 24, and 48 hours after DT washout.

Three biological replicates were obtained. RNA extraction was performed as described previously (see RNA extraction and real-time PCR section). RNA samples were prepared at a concentration of 5 ng/µL in Milli-Q water and submitted to the Utrecht Sequencing Facility (USEQ; UMC Utrecht, The Netherlands) for downstream processing.

Quality control on the sequence reads from the raw FASTQ files was done with FastQC^60^ (v0.11.9). TrimGalore^61^ (v0.6.7) was used to trim reads based on quality and adapter presence after which FastQC was again used to check the resulting quality. rRNA reads were filtered out using SortMeRNA^62^ (v4.3.6) after which the resulting reads were aligned to the reference genome fasta (Mm_GRCm38_gatk_sorted.fasta) using the STAR^63^ (v2.7.10b) aligner. Follow up QC on the mapped (bam) files was done using Sambamba^64^ (v0.8.2), RSeQC^65^ (v5.0.1) and PreSeq^66^ (v3.2.0). Read counts were then generated using the Subread FeatureCounts module^67^ (v2.0.3) with the Mus_musculus.GRCm38.70.gtf gtf file as annotation. Count normalization and log transformation was performed using DESeq2^68^ (v1.48.2). Differential gene expression analysis across all samples was performed using LRT test (DESeq2, reduced model = ∼1). Duplicated genes were excluded from the analysis. Gene set scores were computed using the GSVA^69^ package (v2.2.1). Signature gene sets were obtained from Serra et al.^37^. In **Figure S4K**, a custom proliferation signature was used, including *Mki67, PcnA, Top2a, Cdk1, Ccnb1, Ccnb2, Ccna2, Ccne1, Ccne2, Cdc20, Cdc45, Cdc6, Cdt1, AurkA, AurkB, Plk1, Bub1, Bub1b, Cenpf, Cempe, Mcm2–Mcm7, Tyms, Rrm1, Rrm2, E2f1, Mybl2, Myc, Tfdp1, Ube2c, Espl1, Kif11, and Kif20a*.

### Statistical analysis

Statistical analysis for image analysis was performed in Graphpad Prism 10 or as indicated in the legend. Statistical analysis for RNA sequencing analysis was performed by different packages in R as stated. Details of statistics and sample sizes are described in the figure legends.

## Declaration of generative AI and AI-assisted technologies in the manuscript preparation process

During the preparation of this work, the authors used ChatGPT cautiously to verify the readability of selected sentences and to assist with grammar correction. The authors carefully reviewed and edited the output as needed and take full responsibility for the content of the published article.

## Notes

### Competing Interest Statement

The authors have declared no competing interest.

